# Variegate expression of Cre recombinase in hematopoietic cells in CD11c-cre transgenic mice

**DOI:** 10.1101/2023.07.06.544279

**Authors:** Claire Murat, Sylvie Guerder

## Abstract

In this study, we performed an in-depth analysis of Cre expression in the widely used CD11c-Cre transgenic mice generated by the group of Boris Reizis. In contrast to previous observation, using the highly sensitive Rosa-26-floxed-tdTomato reporter mouse line, we show variegated expression of Cre in multiple hematopoietic linage cells starting in hematopoietic stem cells. We found that in the CD11c-Cre driver mice: (1) Cre is expressed in cDC linage cells and pDC starting from the myeloid dendritic cell precursor, as expected ; (2) Cre is expressed in a substantial fraction of hematopoietic stem cells and common lymphoid progenitors ; (3) Cre is expressed in more than 50% of all leukocytes. Hence, this study indicates that the reporter mice used to characterize Cre expression in Cre-driver mice should be selected with caution and considering the sensitivity of the reporter system. This study also suggests that the interpretation of some reports may need to be re-considered based on a careful evaluation of the cell type-specificity of Cre-mediated in their model.

## Main text

Mice genetically engineered to express Cre recombinase under the control of a tissue- or cell type-specific promoter are powerful tools to analyze the function of a gene of interest in the target tissue or cells and, when crossed with a fluorescent reporter strain, to perform fate-mapping experiments. To generate Cre diver lines, the *cre* transgene is integrated into the marker gene by targeted, knock-in transgenesis. Alternatively, *cre* can be engineered into a bacterial artificial chromosome (BAC) encompassing most, if not all, the transcriptional determinants of the marker gene, which is then randomly inserted into the genome by transgenesis. In these genetically modified mice, Cre expression is expected to recapitulate the physiological expression pattern of the marker gene. In practice, however, the chromatin organization of the locus, and cis-regulatory sequences within the promoter, introns or at distance within the genome may lead to inappropriate spatio-temporal expression of the Cre (1-4).

To confirm tissue-specific expression, Cre driver mice are usually crossed with a strain carrying a gene cassette encoding a fluorescent protein, inserted into the ubiquitously expressed Rosa26 locus. The fluorescent reporter protein will be expressed upon Cre-mediated removal of the *loxP*-flanked stop sequence built into the cassette. Rosa26-floxed-EGFP and Rosa26-floxed-EYFP mice have been widely used to characterized *cre*-transgenic mice. The EGFP and EYPF reporter proteins, however, do not fluoresce brightly and thus lack sensitivity (5, 6). The development of red-fluorescent proteins such as the tdTomato, which is far brighter than EGFP, combined with the introduction, in the Rosa26 locus, of strong promoter/enhancer sequences has improved the sensitivity of Rosa26-reporter mice (7).

To tag all conventional dendritic cells (cDC) and plasmatoid dendritic cells (pDC), the group of Boris Reizis generated a BAC-transgenic mice with the *cre* gene integrated within 160-Kb of the mouse cDC11c gene (CD11c-Cre, (8), B6.Cg-Tg(Itgax-cre)1-1Reiz/J). When crossed with Rosa26-floxed-EYFP reporter mice, Cre-mediated recombination, as evidenced by EYFP expression, was largely confined to the cDC linage with 96% of cDC and 86% of pDC expressing EYFP and minimal recombination in other hematopoietic cells (6% in T cells, 5% in B cells and 16% in NK cells, (8)). To gain in sensitivity we crossed the CD11c-Cre mice with mice expressing the tdtomato in the Rosa26 locus (Rosa26-floxed-tdTomato, B6.Cg-*Gt(ROSA)26Sor*^*tm9(CAG-tdTomato)Hze*^/J) and found variegated expression of the tdTomato in multiple hematopoietic cells. We first analyzed expression in different spleen cell subsets (staining strategy in Sup. Fig 1).

As expected we found that all cDC expressed high level of tdTomato starting at the precursor stage (pre-cDC) as well as all pDC (Fig 1 A-B). Unexpectedly, we found that a large fraction of CD4 and CD8 T cells and B cells from spleens also expressed high levels of tdTomato indicating that the Cre recombinase is also expressed in these cells (Fig 1 C-D). A fraction of NK cells also express tdTomato and this was not confined to the CD11c positive NK cells (Fig. 1 C-D and not shown). This contrast with the phenotype of spleen cells in the CD11c-DTR-EGFP mice in which GFP expression is confined to the cDC1 and cDC2 subsets (Fig. 1E-F).

**Figure 1:**
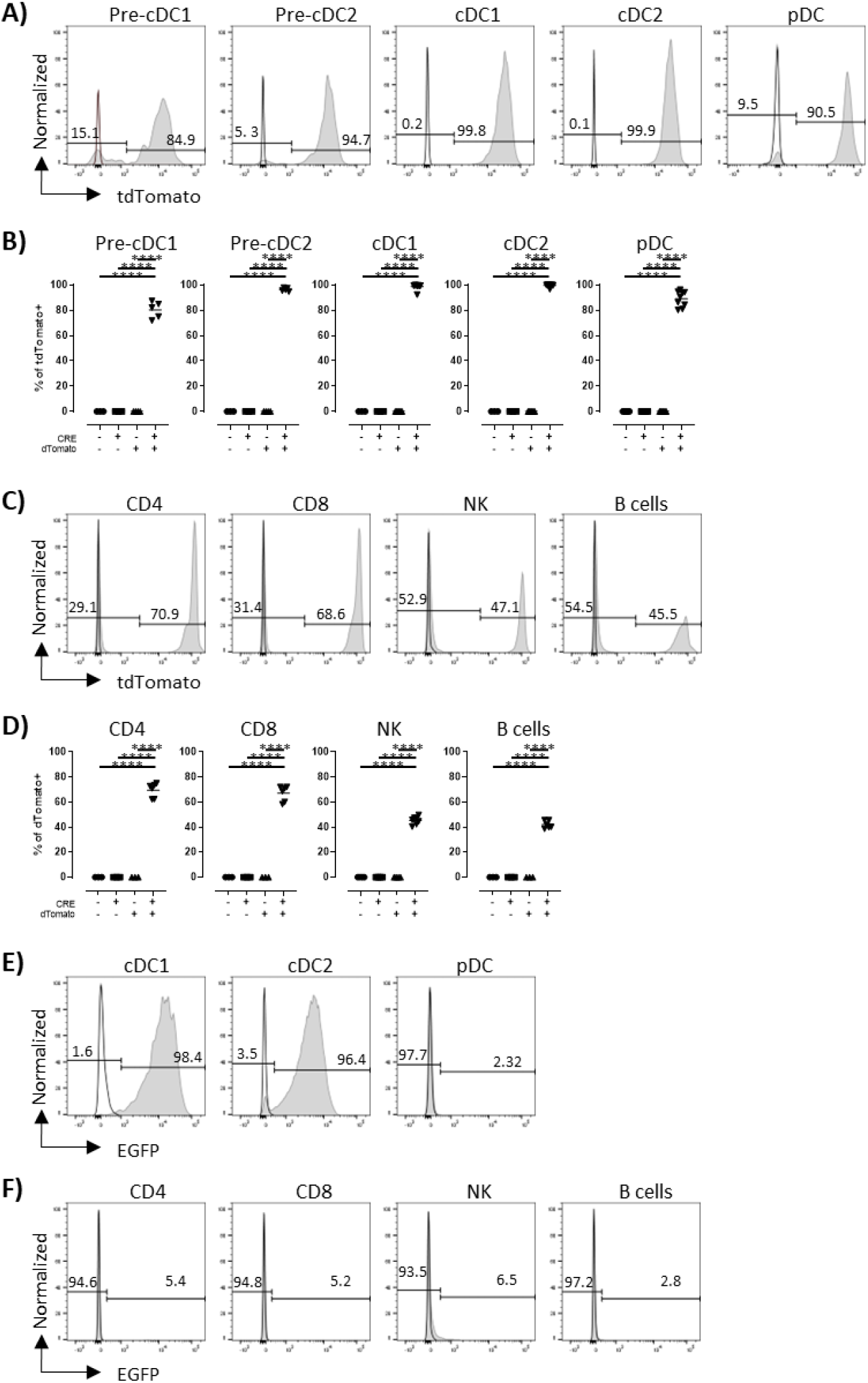
tdTomato expression into myeloid and lymphoid populations from the spleen in CD11c-CRE-tdTomato mice. **A-D)** Expression of tdTomato in spleenocytes is shown. **A, C)** Representation of the single cell expression of tdTomato for the indicated spleen cell population from the CD11c-Cre-tdTomato double transgenic (filled grey histogram) or negative littermate (black line) **B, D)** Percentage of the indicated cells expressing tdTomato in CD11c-Cre x Rosa26-floxed-tdTomato progeny according to their genotype. **E-F)** Representation of the single cell expression of EGFP for the indicated spleen cell population from the CD11c-DTR-EGFP transgenic mice (filled grey histogram) or negative littermate (black line). Each symbol corresponds to one mouse analyzed in 2 to 5 independent experiments. Unpaired Student t-tests test are shown as * P < 0.05, ** P < 0.01, ***P<0,001 and ****P<0,0001.

To define the developmental stage of Cre expression in CD11c-Cre mice we performed an in-depth analysis of their hematopoietic precursors. Bone marrow cells (BM) were fractionated by FACS staining as describe in sup Fig. 2. Low levels of tdTomato expression was detected in 27.2±2.3% (mean+SD) of the most early hematopoietic precursors, the hematopoietic stem cells (HSP) and multipotent progenitor (MMP, Fig. 2A-B). Expression of tdTomato was also detectable thought in a very small fraction of granulocyte and monocyte precursors (GMP/cMoP) and monocytes of the BM (Fig. 2 C-D). In the DC lineage, expression of tdTomato started in the myeloid dendritic cell (MDP) precursor with 12.1±1.1% of MDP being tdTomato positive and increased thereafter throughout cDC differentiation with 29.6±1.1% of the common dendritic cell precursor (CDP) and 79.4±2.8 and 92.8±1.2% of the pre-cDC1 and pre-cDC2, respectively, expressing tdTomato (Fig. 2 E-F). Hence, within the granulocyte and myeloid lineage Cre-mediated recombination mainly start when precursors committed to the DC lineage.

**Figure 2:**
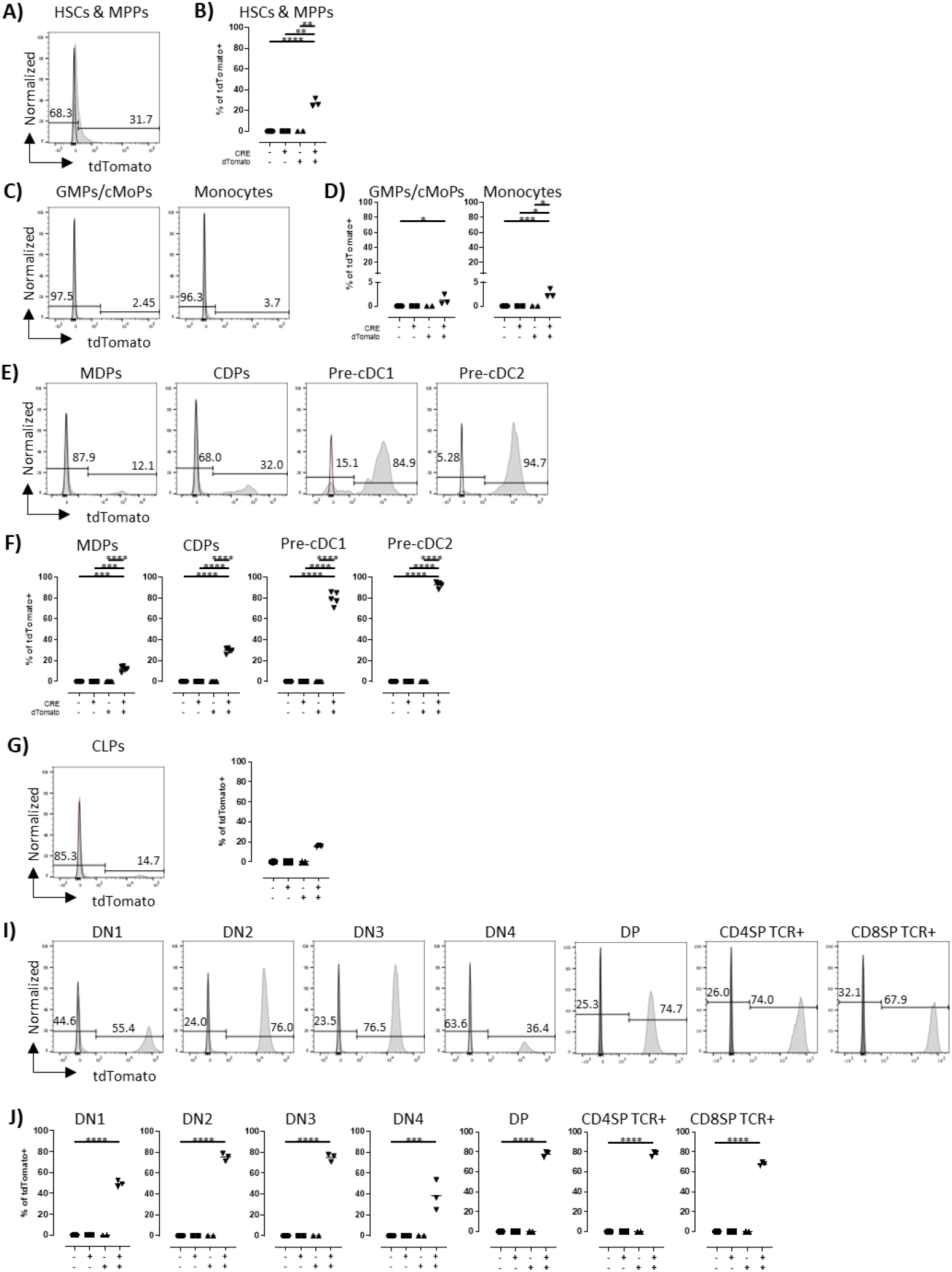
tdTomato expression in lymphoid and myeloid hematopoietic precursors in CD11c-CRE-tdTomato mice. **A, C, E, G, I)** Representation of the single cell expression of tdTomato in the indicated BM cell population from CD11c-Cre-tdTomato double transgenic mice (filled grey histogram) or negative littermate (black line). **B, D, F, H, J)** Percentage of the indicated BM precursors expressing tdTomato into CD11c-Cre x Rosa26-floxed-tdTomato progeny according to their genotype. Each symbol corresponds to one mouse analyzed in 2 to 3 independent experiments. Unpaired Student t-tests test are shown as * P < 0.05, ** P < 0.01, ***P<0,001 and ****P<0,0001.

Within the lymphoid lineage, 16.6±0.2% of common lymphoid precursors (CLP) expressed tdTomato at high levels indicating that Cre is expressed at significant levels in early lymphoid precursors (Fig 2 G-H). Regarding T cell differentiation in the thymus, double negative cells (DN) expressed tdTomato at increasing frequency throughout differentiation from DN1 to DN3 (Fig 2 I-J). The frequency of tdTomato positive DN4 cells was lower as compared to DN2-DN3 cells which may be linked to the transcriptional changes associated with β-selection. At the double positive stage, the frequency of tdTomato positive cells raised again to level similar to that of the DN3 and was maintained at similar levels in thymic and peripheral CD4 and CD8 T cells (Fig. 2 I-J). Altogether these results show that in CD11c-Cre transgenic mice, Cre is expressed in all lymphoid cells starting at the CLP stage.

By using a highly sensitive reporter mouse line, we provide here compelling evidence of Cre expression in different hematopoietic lineage cells in the CD11c-Cre transgenic mice. This expression does not merely reflect transient CD11c expression during hematopoiesis as indicated by scRNAseq analysis (Immgen) but instead leaky expression of the transgene. Testing Cre transgenic mice using highly sensitive fluorescent report line is therefore essential to carefully control the expression pattern of the Cre recombinase. Yet, given that chromatin accessibility could constrain Cre-mediated recombination, leaky expression of Cre might not necessary lead to inappropriate recombination in unwanted cell type, something that need to be controlled by additional approaches.

## Acknowledgments

We thank the SG’s lab for helpful discussion and manuscript edition. We thank the personnel of ANEXPLO/US006 for animal husbandry; Fatima-Ezzahra L’Faqihi, Valérie Duplan and Anne-Laure Iscache for technical assistance at the Flow cytometry facility TRI-CPTP, Toulouse;

## Author Contribution

S.G designed the experiments and wrote the manuscript with contribution from C.M. who performed experiments.

## Material and methods

### Mice

CD11c-Cre mice (B6.Cg-Tg(Itgax-cre)1-1Reiz/J) were crossed with Rosa26-floxed-tdTomato mice (B6.Cg-*Gt(ROSA)26Sor*^*tm9(CAG-tdTomato)Hze*^/J) to generate CD11c-Cre-tdTomato double transgenic, CD11c-Cre and Rosa26-floxed-tdTomato single transgenic mice and non-transgenic littermate. Experimental mice were used at 6-25 weeks of age. All mice were housed under specific and opportunistic pathogen-free conditions at US006, Toulouse and experiments were performed in accordance with National and European regulations and guidelines. Mouse experimental protocols were approved by the French ‘Ministère de l’Enseignement Supérieur et de la Recherche’ (APAFIS#25920-202005261836395 v5)

### Cells isolation and FACS staining

Spleens or thymi were first dissociated with 125 μg/ml Liberase TL (Roche) and 40 μg/ml DNAse I (Sigma) in RPMI-2% FCS for 20 min at 37°C. Bone marrow (BM) cells were flushed out of the tibias and femurs. Then, erythrocytes were removed using Gey’s treatment. After 2 washes, cells were incubated in 2.4G2 for 30 min and surface stained using standard procedure. The following antibodies were obtained from eBioscience, BD Pharmingen or Miltenyi: anti-B220-FITC, PerCP-Cy5.5 or BV786 (RA3-6B2), anti-CD115-Pe-Vio770 (REA827), anti-CD11c-APC-H7, BUV395 or Pe-Cy7 (HL3 or N418), anti-CD127-APC (SB/199), anti-CD135-Biotin (A2F10), anti-CD19-BV480 (1D3), anti-CD24-BV421 (M1/69), anti-CD25-BV421 (PC61), anti-CD34-FITC (RAM34), anti-CD4-Pacific blue, Pe-Cy7 or APC, anti-CD44-BV786 (IM7), anti-CD8-APC or PerCP-Cy5.5 (53-6.7), anti-CD90.2-BV510 (53-2.1), anti-MHC-II-BV605, BV711 or PerCP-Vio700 (M5/114.15.2), anti-CD49b-Pacific Blue or FITC (DX5), anti-Nkp46-BV510 (29A1.4), anti-Sirpα-BV711 (P84), anti-TCRβ-FITC (H57597) and streptavidin Pe-Cy7, APC or PE. Dead cells were excluded using Fixable viability dye EF506 (eBioscience). Data were collected on BD Symphony A5 (BD Biosciences) cytometers and analyzed using FlowJo software (Tree Star).

### Statistical Analysis

Mean values and unpaired student t-test were calculated with GraphPad Prism (GraphPad software).

## References

1. Becher B, Waisman A, and Lu LF. Cre-lox: Target Sensitivity Matters. Immunity. 2019.51:595.

2. Reizis B. The Specificity of Conditional Gene Targeting: A Case for Cre Reporters. Immunity. 2019. 51:593–594.

3. Schmidt-Supprian M, and Rajewsky K. Vagaries of conditional gene targeting. Nat Immunol. 2007. 8:665–668.

4. Song AJ, and Palmiter RD. Detecting and Avoiding Problems When Using the Cre-lox System. Trends Genet. 2018. 34:333–340.

5. Mao X, Fujiwara Y, Chapdelaine A, Yang H, and Orkin SH. Activation of EGFP expression by Cre-mediated excision in a new ROSA26 reporter mouse strain. Blood. 2001. 97:324–326.

6. Srinivas S, Watanabe T, Lin CS, William CM, Tanabe Y, Jessell TM, et al. Cre reporter strains produced by targeted insertion of EYFP and ECFP into the ROSA26 locus. BMC Dev Biol. 2001. 1:4.

7. Dou Y, Lin Y, Wang TY, Wang XY, Jia YL, and Zhao CP. The CAG promoter maintains high-level transgene expression in HEK293 cells. FEBS Open Bio. 2021. 11:95–104.

8. Caton ML, Smith-Raska MR, and Reizis B. Notch-RBP-J signaling controls the homeostasis of CD8-dendritic cells in the spleen. J Exp Med. 2007. 204:1653–1664.

