## Supplementary figures for "Variegate expression of Cre recombinase in hematopoietic cells in CD11c-cre transgenic mice"

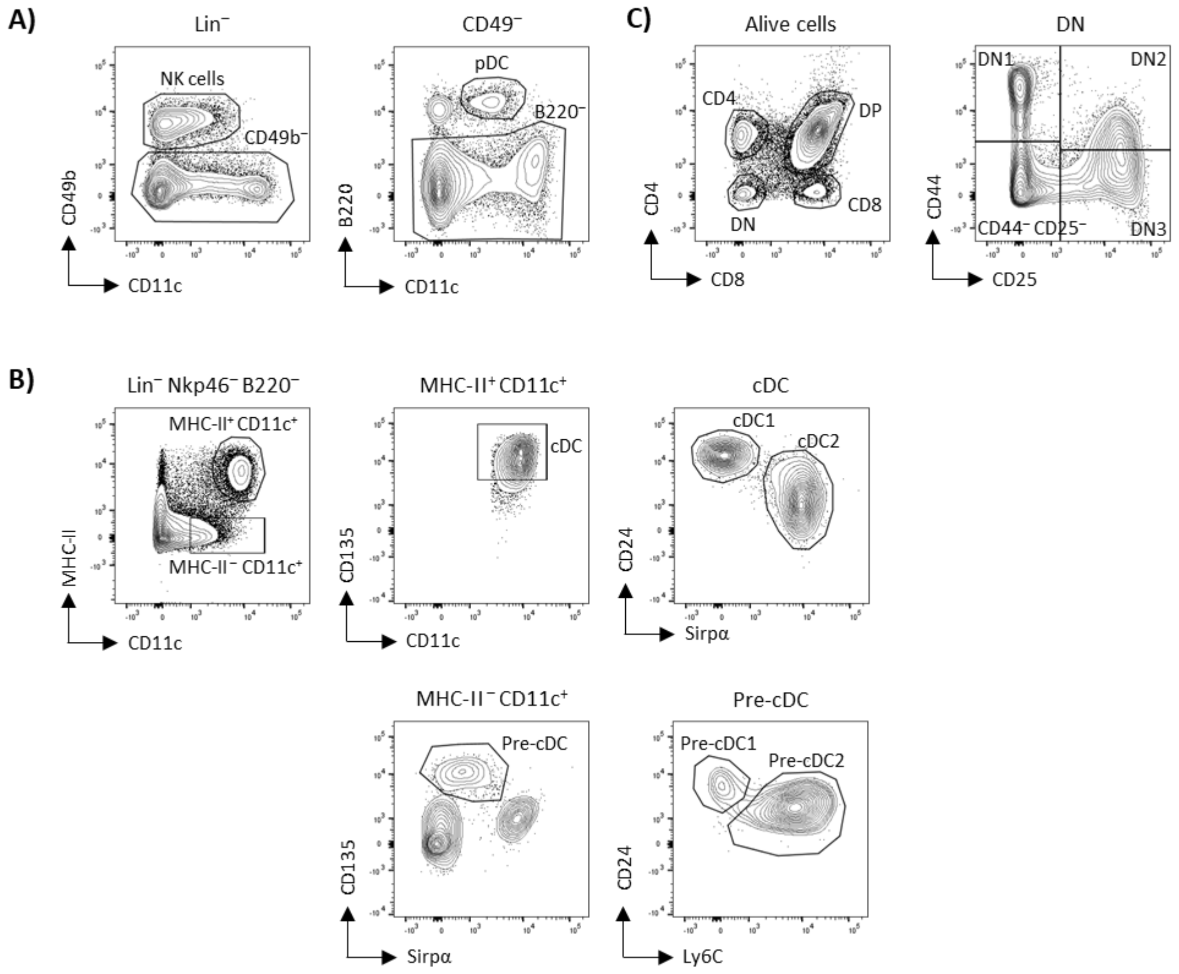


**Supplementary Fig. 1: Flow cytometry characterization of the different populations of interest in the spleen and thymus.**

**A)** Cells were stained with PO- conjugated anti-CD90, -CD19 and -Ly6G antibodies and Fixable viability dye EF506 to exclude T cells, B cells, polymorphonuclear and dead cells (Lin$^{-}$). Among Lin$^{-}$ cells, NK cells were identified as CD49b$^{+}$ CD11c$^{int}$ (left panel). Among Lin$^{-}$CD49b$^{-}$ cells, pDC were identified as B220$^{+}$ CD11c$^{int}$ (right panel). **B)** Among Lin$^{-}$ B220$^{-}$ Nkp46$^{-}$ cells cDC were identified as the MHC-II$^{+}$ CD11c$^{+}$ CD135$^{+}$ cells and cDC1 and cDC2 subsets based on CD24 and Sirpα expression, respectively (right, upper panel). The pre-cDC were identified as MCH-II$^{-}$ CD11c$^{+}$ CD135$^{+}$ Sirpα$^{-}$ cells and the pre-cDC1 and pre-cDC2 subsets were identified based on CD24 and Ly6C, respectively (right, lower panel). **C**) Live cells are plotted for CD4 and CD8 to define Double Positive (DP), CD4$^{+}$ T, CD8$^{+}$ T cells and Double Negative (DN) cells as indicated. Among the DN cells, the DN1, DN2 and DN3 cells were identified based on CD44 and C25 expression as indicated.

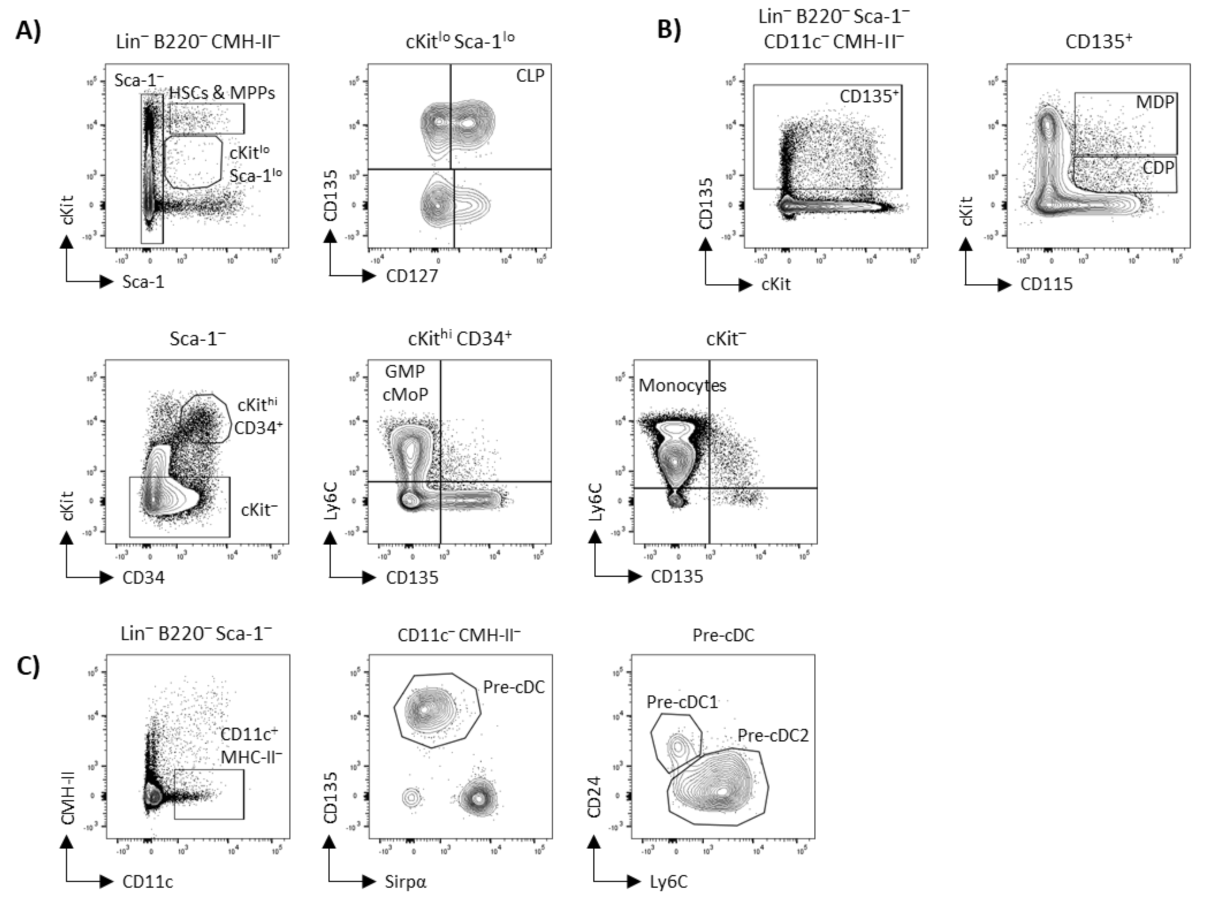


**Supplementary Fig. 2: Flow cytometry characterization of the different hematopoietic progenitors in the bone marrow.**

**A)** Cells were stained for CD3 (Lin$^-$), B220, MHC-II markers and Fixable viability dye EF506 to exclude T cells, B cells, DC and dead cells. Among Lin$^-$ B220$^-$ MHCII$^-$ cells, hematopoietic stem cells (HSCs) and multipotent progenitors (MPPs) were identified as cKit$^{hi}$ Sca-1$^{hi}$ cells (upper, left panel). Among the cKit$^{lo}$ Sca-1$^{lo}$ cells, Common Lymphoid Progenitors (CLPs) we identified based on the expression of CD135 and CD127 (upper, right panel). Within the Sca-1$^-$ population, the cKit$^{hi}$ CD34$^+$ cells include the Granulocyte Macrophage Progenitor (GMPs) and Common Monocyte Progenitors (cMoPs) characterized by expression Ly6C and the cKit$^-$ cells include monocytes which are identified based on Ly6C expression. **B)** Cells were stained with for CD3, Nkp46, Ly6G and Fixable viability dye EF506 to exclude T cells, NK cells, polymorphonuclear and dead cells (Lin$^-$). Among Lin- cells, B220, Sca-1, CD11c and MHC-II markers were used to exclude B cells, hematopoietic progenitors and cDC, then Common Myeloid Progenitors (CMPs) and Common Dendritic cell Progenitors (CDPs) were identified as: CD135$^+$ CD115$^+$ cKit$^{hi}$ and CD135$^+$ CD115$^+$ cKit$^{int}$, respectively. **C)** Cells were stained for CD3,

Nkp46, Ly6G and Fixable viability dye EF506 to exclude T cells, NK cells, polymorphonuclear and dead cells (Lin$^{-}$). Among Lin- cells**,** B220 and Sca-1 markers were used to exclude B cells, pDC and hematopoietic progenitors then CD11c$^{+}$ MHC-II$^{-}$ cells were gated. Within the CD11c$^{+}$ MHC-II$^{-}$ population, pre-cDC were identified as CD135$^{+}$ Sirpα$^{-}$ (middle panel) and pre-cDC1 and pre-cDC2 subset were then defined according to their expression of CD24 and Ly6C, respectively (right panel) .
